# A PBD-dimer containing antibody drug conjugate targeting CCRL2 for high-risk MDS/AML

**DOI:** 10.1101/2025.08.17.670714

**Authors:** Nour Sabiha Naji, Taha Ahmedna, J. David Peske, Xinghan Zeng, Brandy Perkins, Zanshe Thompson, Tushar D. Nichakawade, Bum Seok Lee, Evangeline Watson, Theodora Chatzilygeroudi, Li Luo, Bogdan Paun, Melanie Klausner, Yuju An, Teodora Supeanu, Ivana Gojo, Gabriel Ghiaur, Amy E. DeZern, Mark J. Levis, Linda Resar, Richard J. Jones, Styliani Karanika, Suman Paul, Theodoros Karantanos

## Abstract

Patients with myelodysplastic syndrome (MDS)/acute myeloid leukemia (AML) with high-risk features including *TP53* mutations and deletions have poor outcomes due to lack of effective therapies. The atypical chemokine surface receptor C-C motif chemokine receptor-like 2 (CCRL2) is overexpressed in MDS and secondary AML (sAML) compared to healthy hematopoietic cells and we recently found that *TP53*-mutated MDS/AML and AML with erythroid features express the highest levels of this receptor across MDS/AML subtypes. To illustrate the therapeutic potential of CCRL2 as a therapeutic target, we developed an anti-CCRL2 antibody-drug conjugate (ADC) by conjugating an anti-CCRL2 antibody with the cytotoxic drug pyrrolobenzodiazepine (PBD), which causes DNA double-strand breaks leading to cancer cell death. The anti-CCRL2 ADC demonstrated strong CCRL2-selective cytotoxicity against cell lines derived from MDS/AML patients with *TP53* mutations and erythroid features, surpassing the cytotoxic effects observed with gemtuzumab and PBD-conjugated anti-CD33 and anti-CD123 ADCs. It also induced apoptosis and suppressed the clonogenicity of primary MDS/AML bone marrow samples without affecting the survival, differentiation and clonogenicity of healthy hematopoietic stem and progenitor cells. This agent also suppressed the leukemic growth of *TP53-*mutated MDS/AML cell line xenografts, improving mice survival and decreasing the leukemic burden in patient-derived *TP53*-mutated MDS/AML xenografts. In conclusion, our study introduces CCRL2 as a potential new therapeutic target in high-risk MDS/AML.

**Statement of significance:** Pyrrolobenzodiazepine(PBD)-conjugated anti-CCRL2 ADC shows anti-leukemic effect in MDS/AML models including *TP53*-mutated disease without affecting healthy hematopoietic cells supporting that it is a promising candidate for single-agent or combination therapies in high-risk MDS/AML.

## Introduction

Patients with high-risk myelodysplastic syndrome (MDS) and secondary acute myeloid leukemia (sAML) continue to exhibit poor outcomes, especially in the presence of high-risk features such as *TP53* mutations or deletions [1–3] Thus, the development of effective therapies is urgently needed for these individuals. Antibody-drug conjugates (ADCs) [4, 5], show prominent activity in lymphoid cancers[6, 7], and good-risk AML [8, 9]. However, they demonstrate limited efficacy in high-risk MDS/AML[10].

The atypical chemokine receptor C-C motif chemokine-like receptor 2 (CCRL2) is normally expressed in differentiated myeloid cells contributing to the regulation of cells migration in inflammation cites [11–13]. This surface receptor is upregulated in MDS and sAML, while normal hematopoietic progenitors demonstrate minimal expression[14]. CCRL2 silencing suppresses MDS/sAML cell growth and sensitizes them to hypomethylating agents but does not affect the survival and clonogenicity of healthy CD34+ cells[14, 15]. We recently discovered that across MDS/AML subtypes, *TP53* mutated MDS/AML and AML with erythroid features express the highest CCRL2 levels[16]. Intriguingly, *TP53* deletion in AML *TP53* wild-type (WT) cells caused a direct prominent CCRL2 upregulation[16]. The main downstream target of CCRL2 is interferon gamma signaling, which has been associated with AML clonal evolution, *TP53* deletion, acquisition of erythroid features and treatment resistance[16–18]. Thus, CCRL2 is potentially a promising target in in high-risk MDS/sAML.

In this study, we show that an ADC targeting CCRL2 exhibits prominent anti-leukemic effects in high-risk MDS/AML, including *TP53*-mutated cells, but has limited toxicity against normal hematopoietic cells.

## Material and Methods

### Cell lines and reagents

TF-1, SKM1, SET2, and MV4-11 cell lines were purchased from the American Type Culture Collection. F36P, OCI-AML3 and MOLM13 cells were purchased from Leibniz Institute DSMZ. MDS-L cells were a gift from Dr. Starczynowski, University of Cincinnati [30]. MV4-11 cells were cultured in IMDM (Thermo Fischer Scientific, Waltham, MA) with 10% fetal bovine serum (FBS) (MilliporeSigma, Burlington, MA). TF-1, MOLM13 and MDS-L were cultured in RPMI 1640 (Thermo Fischer Scientific, Waltham, MA) with 10% FBS and F36P, SET2 and SKM-1 with 20% FBS with the addition of GM-CSF for TF-1 and F36P (2 ng/ml and 20 ng/ml respectively; PeproTech). All the cell lines were cultured with 2mM L-glutamine, penicillin (100 U/ml) and streptomycin (100µg/ml) at 37 in 5% CO2. Doxycycline was purchased from Sigma Aldrich (D9891) and was diluted in PBS (Thermo Fischer Scientific, Waltham, MA). Gemtuzumab (mylotarg) was purchased from MedChem Express (#HY-109539).

### ADCs generation

The anti-CD33/CD123/humanCCRL2/mouseCCRL2 and isotope controls (IgG2a and IgG2b) ADCs were developed by conjugating the anti-CD33 (Santa Cruz #374450), anti-CD123 (BD Biosciences #554527), anti-humanCCRL2 antibody (BioLegend #358302), anti-mouseCCRL2 antibody (BioLegend #114002), IgG2a and IgG2b isotype control antibodies (BioLegend # 400502/ 400602) with PBD using a dipeptide linker (SG3249 (MedChemExpress Cat. No.: HY-128952). was done by reducing monoclonal antibodies using 5x molar excess Tris(2-carboxyethyl) phosphine hydrochloride (TCEP, ThermoFisher Scientific, Cat. No: 20490) followed by TCEP removal using Zeba Spin columns (ThermoFisher Scientific, Cat No: 89882). Subsequently, antibodies were reacted with 5x molar excess SG3249 in 10% DMSO at room temperature. Excess unreacted drug linkers were subsequently removed by buffer exchange into PBS using Zeba Spin columns. ADC drug conjugation was analyzed by hydrophobic interaction chromatography (HIC) using Agilent 1260 Infinity I LC system as previously described. Baseline correction and analysis of HPLC spectra was performed using the OriginPro v10.1.5.132. The HPLC methods and drug-antibody ratio (DAR) calculations were described before[19].

### MTT assay

TF-1, F36P, K562, THP1, SET2, MOLM13, MV4-11, MDS-L and SKM1 cells were plated in 96-well plates (7,500 cells per well) and treated for 6 days with anti-CCRL2, anti-CD33, anti-CD123, gemtuzumab and IgG2a antibodies. The 3-(4,5-dimethylthiazol-2-yl)-2,5-diphenyl-2*H*-tetrazolium bromide (MTT) assay (11465007001, Roche Diagnostics, Mannheim, Germany) was conducted according to the manufacturer’s instructions. Absorbance was measured at 570 nm. Calculation of cell viability by 100x (absorbance of sample/average absorbance of untreated control) for the respective dose for each cell line.

### Flow cytometry analysis

Healthy control (HC) samples were stained with FITC-conjugated anti-human CD34 (BioLegend, #343603), BV605-conjugated anti-human CD38 (BioLegend, #303531), BV421-conjugated anti-human CD71(Biolegend, #334121), and APC-conjugated anti-human CD235a (#349113). Cell lines were stained with PE-conjugated anti-CCRL2 (BioLegend, #358303), APC-conjugated anti-CD123 (BioLegend #973704), and PE-conjugated anti-CD33 (BioLegend #303403). Primary samples were stained with BV510-conjugated anti-human CD45 (BioLegend, #103137) and PE-conjugated anti-CCRL2. Cell line and patient-derived xenograft samples were stained with PE-Cy7-conjugated anti-mouse CD45 (BioLegend #103113), and BV510-conjugated anti-human CD45 (BioLegend, #103137). Gating was based on an unstained control. For the assessment of apoptosis, cells were stained with 7AAD (BioLegend, #420403). Following staining, analysis was performed using BD LSR II (BD Biosciences). Mean fluorescence intensity (MFI) was measured for each marker using FlowJo analysis software version 10.0.8(FlowJo, Ashland, CO, USA).

### CCRL2 knockout (KO)

Lentiviral vectors expressing CCRL2-targeting sgRNA (pLV [CRISPR]-hCas9:T2A: Puro-U 6>hCCRL2[gRNA#162], pLV [CRISPR]-hCas9:T2A: Puro-U6>hCCRL2 [gRNA#177]) or empty pLKO.1-puro lentiviral vector (pLV [CRISPR]-hCas9/Puro-U6>Scramble_gRNA1) was transfected into 293T cells using Lipofectamine 2000 (Thermo Fischer Scientific) for lentiviral supernatant production.TF-1, F36P and SET2 cells were incubated with the viral supernatant and polybrene (8µg/ml; MilliporeSigma) for transduction. 48 hours later, cells were treated with puromycin (2µg/ml for TF-1, 1.5 µg/ml for F36P and 0.75 µg/ml for SET2) for 3-4 days for resistant cells selection.

### Colony formation assay

Clonogenic assays were conducted as previously detailed [15, 16]. Cells were counted and resuspended at a density of 3000 cells/ml in methylcellulose-based media. Following around two weeks of incubation at 37°C in 5% CO2, counting of colony forming units was performed under bright-field microscopy. A colony was defined as a cell aggregate of >50 cells.

### Patients and sample processing

All specimens were obtained by the Johns Hopkins Kimmel Cancer Center Specimen Accessioning Core. In accordance with the Declaration of Helsinki and under a research protocol approved by the Johns Hopkins Institutional Review Board, informed consent was procured from all donors before specimen collection. Isolation of CD34+ cell subsets was performed using the CD34 MicroBead kit (Miltenyi Biotec) as before. Samples were collected from marrow aspirations of multi-hit *TP53*-mutated and wild-type MDS/AML or de novo AML patients. Control marrow was collected as excess material after harvesting healthy donors for allogeneic bone marrow transplantation. ADCs (20 or 40 ng/ml for *TP53*-mutated or *TP53* wild-type MDS/AML primary cells and 100 ng/ml for HC MNCs or CD34+ cells) were added at day 0 to MNCs or sorted CD34+ cells. At day 5, apoptosis was assessed by 7AAD staining and, differentiation was assessed by CD34, CD38, CD71, CD235a staining for HC samples. Clonogenicity was assessed by plating 2,000 cells in methylcellulose-based medium (StemCell #04434).

### Treatment of C57BL/6 mice

Total 14 8-10 weeks old C57BL/6 mice were treated intravenously with 1 mg/kg anti-mouse CCRL2 ADC or conjugated IgG2b. Complete blood counts and chemistry panels were analyzed before treatment and 4 weeks later. Weight was measured every week. Mice were harvested at 4 weeks post-treatment and kidney and spleen weights were measured.

### Cell line xenografts

Luciferase+ TF-1, SKM1 and F36P cells were injected to 8-10-week-old NOD.Cg-*Prkdc^scid^ Il2rg^tm1Wjl^*/SzJ (NSG) female mice (10^6^ cells per mouse) [14, 15]. Using IVIS spectrum *in vivo* imaging system, bioluminescence signal was measured at different timepoints. At day 10 or 15, mice were randomized to receive either conjugated IgG2a or anti-CCRL2 ADC (0.75 mg/kg for TF-1, 1 mg/kg for SKM1 and F36P, intravenously). Among F36P engrafted mice, 5 vs 5 mice were sacrificed on day 35 (after one dose of treatment) to assess disease burden and 4 vs 4 mice were monitored until day 65. Survival of the mice was assessed by day 65 for TF-1, 50 for SKM1 and 75 for F36P. At that time, all remaining mice were euthanized, and the percentage of human CD45+ cells was assessed in bone marrow, spleens or livers by flow cytometry. IACUC guidelines were followed.

### Patient-derived multi-hit *TP53-*mutated MDS/AML xenografts

MNCs collected from marrow aspirates of multi-hit *TP53*-mutated MDS/AML were injected to sub-lethally irradiated (1.5 Gy) 8-10-week-old NOD.Cg-*Prkdc^scid^ Il2rg^tm1Wjl^* Tg (CMV-IL3, CSF2, KITLG)1Eav/MloySzJ (NSGS) female mice (5×10^5^ cells/mouse). Bone marrow aspirates and peripheral blood were collected from mice at day 30 and based on disease burden, engrafted mice (≥2% human CD45+ cells in bone marrow) were randomized to receive conjugated IgG2a or anti-CCRL2 ADC (1 mg/kg). At day 55, peripheral blood, spleens, livers and bone marrows were collected from mice to assess disease burden by flow cytometry and immunohistochemical analysis. IACUC guidelines were followed.

### Immunohistochemistry

Following preparation of paraffin bone marrow, spleen and liver sections, slides were prepared and stained using hematoxylin and eosin (H&E).

### Statistical Analysis

Statistical analysis was performed by using GraphPad Prism (GraphPad Software, La Jolla, CA). Mann-Whitney test was performed to assess statistical significance when comparing two groups. One-way analysis of variance (ANOVA) was performed for the comparisons of three or more groups. Kaplan-Meier analysis and mantle Cox log rank test was performed for comparisons of mice survival. Standard deviation was used to assess centrality and dispersion.

## Results

### Anti-CCRL2 ADC exhibits prominent cytotoxicity against MDS/AML cell lines

To identify the best payload for the development of an ADC with high activity against high-risk MDS/AML, five commonly used ADC payloads (DM1, MMAE, Exatecan, SN38, PBD) were compared and pyrrolobenzodiazepine (PBD) showed the highest cytotoxicity against the erythroleukemic/*TP53*-mutated TF-1 cells, which was not affected by CCRL2 expression based on the comparison of CCRL2 wild-type (WT) or knock-out (KO) cells (**Supplementary1A-B**). The (PBD) dimer (SG3199)[20]-conjugated ADC loncastuximab has received FDA approval for high-grade B-cell lymphomas treatment[21, 22]. Thus, an anti-CCRL2 ADC was developed by conjugating a monoclonal anti-human CCRL2 antibody, with PBD. A PBD-conjugated IgG2a antibody (IgG2a ADC) was used as control. Conjugations were confirmed by high-performance liquid chromatography (**Supplementary Figurebold>, D**) showing a drug-antibody ratio (DAR) of ∼3.7 (**Supplementary Table 1**).

To assess anti-CCRL2 ADC selective efficacy against cells expressing CCRL2, CCRL2 WT and KO TF-1, F36P and SET2 [16] cells, were treated with increasing doses (0-300 ng/ml) of ADCs for 6 days. The anti-CCRL2 ADC had a significantly higher cytotoxicity against WT compared to KO cells and compared to conjugated IgG2a (**Figure 1A-C**). To further assess CCRL2-selective activity doxy-inducible CCRL2 TF-1 cells (iTF-1) which are genetically engineered TF-1 cells to express CCRL2 under doxycycline [15], were used. The activity of anti-CCRL2 ADC was significantly higher when cells were treated with doxycycline compared to untreated ones (**Figure 1D**).

**Figure 1.**
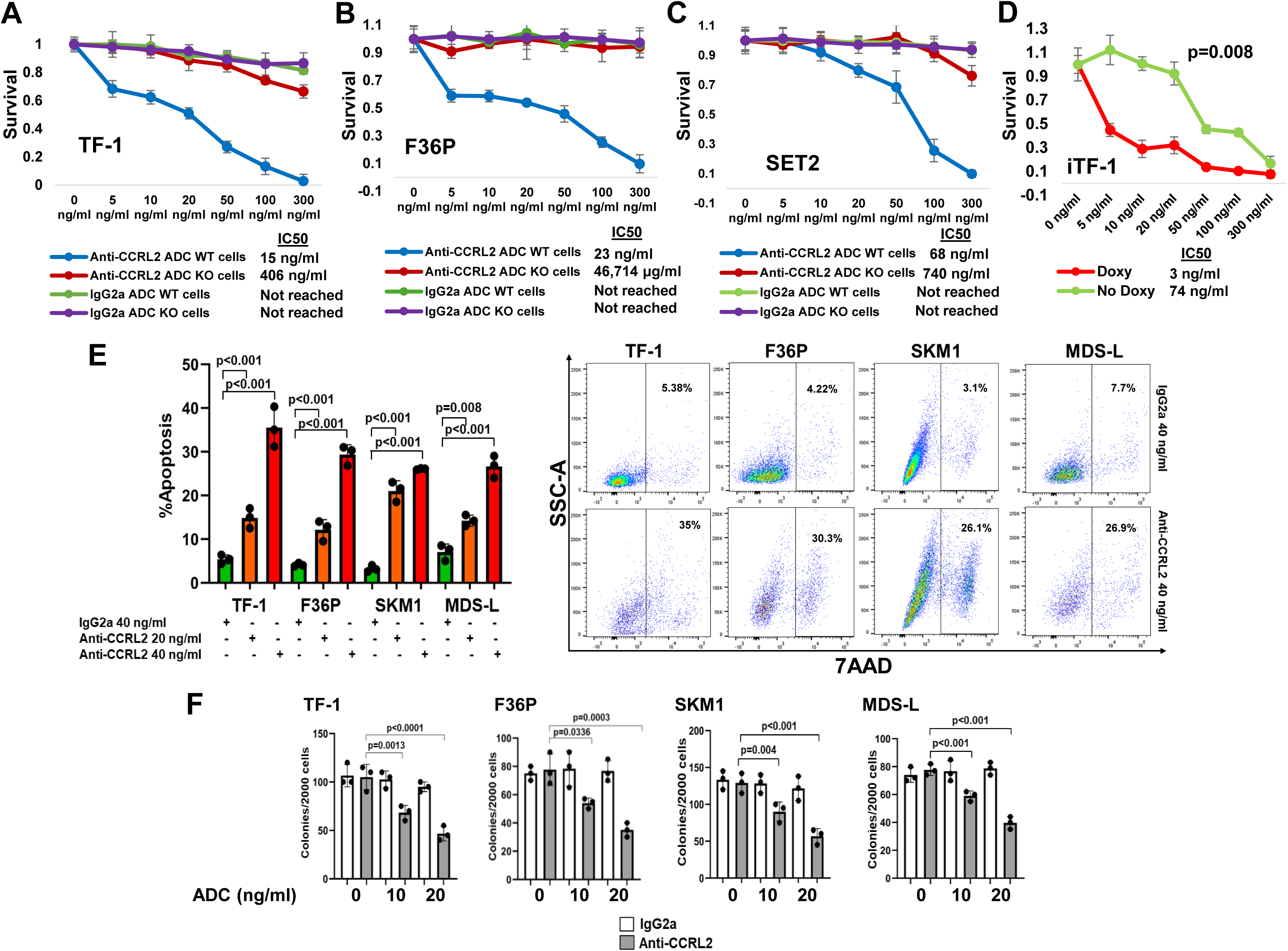
Anti-CCRL2 ADC exhibits prominent cytotoxicity against MDS/AML cell lines. **(A - C)** CCRL2 wildtype (WT) and knockout (KO) TF-1, F36P and SET2 cells were treated with increasing doses of the ADCs (0 ng/ml,5 ng/ml,10 ng/ml, 20 ng/ml, 50 ng/ml, 100 ng/ml and 300 ng/ml) for 6 days and showed significantly higher toxicity in TF1 WT (IC50=15 ng/ml, p<0.001), F36P WT (IC50=23 ng/ml, p<0.001) and SET2 WT (IC50=68 ng/ml, p<0.001) cells treated with the anti-CCRL2 ADC compared to KO cells (IC50=406 ng/ml for TF-1 KO, IC50= 46,714 ng/ml for F36P KO and IC50=740 ng/ml for SET2 KO) or cells treated with the conjugated IgG2a. **(D**) Doxy-inducible CCRL2 TF-1 cells were treated with increasing doses of anti-CCRL2 ADC in the presence or absence of 10 ng/ml doxycycline showing significantly higher cytotoxicity in cells treated with doxycycline (IC50 for doxy-treated=3 ng/ml vs. IC50 for untreated= 74 ng/ml, p=0.008). **(E)** TF-1, F36P, SKM1 and MDS-L cells were treated with 20 or 40 ng/ml of anti-CCRL2 ADC or 40 ng/ml anti-IgG2a ADC. Apoptosis was measured by flow cytometry using 7AAD stain. Anti-CCRL2 ADC induced apoptosis in both TF1 (p=0.002 for 20 ng/ml and p<0.001 for 40 ng/ml), F36P (p=0.003 for 20 ng/ml and p<0.001 for 40 ng/ml), SKM1 (p<0.001 for 20 ng/ml and 40 ng/ml) and MDS-L (p=0.008 for 20 ng/ml and p<0.001 for 40 ng/ml) cells compared to anti-IgG2a ADC after 6 days of treatment. **(F)** TF1, F36P, SKM1 and MDS-L cells were treated with 10 and 20 ng/ml of anti-CCRL2 ADC. Anti-CCRL2 ADC significantly suppressed clonogenicity of TF-1(p=0.0013 for 10 ng/ml and p<0.0001 for 20 ng/ml), F36P (p=0.0336 for 10 ng/ml and p=0.0003 for 20 ng/ml), SKM1 (p=0.004 for 10 ng/ml and p<0.001 for 20 ng/ml) and MDS-L( p<0.001 for 10 ng/ml and 20 ng/ml) cells.

Treatment with 20 or 40 ng/ml anti-CCRL2 ADC induced apoptosis in TF-1, F36P, SKM1 and MDS-L cells, which all resemble AML with erythroid features (TF-1 and F36P) and MDS/AML with loss-of-function *TP53* mutations (SKM1 and MDS-L cells) compared to treatment with 40 ng/ml conjugated IgG2a (**Figure 1E**). Similarly, 10 and 20 ng/ml anti-CCRL2 ADC suppressed clonogenicity in these cells compared to treatment with the same doses of conjugated IgG2a (**Figure 1F**).

Collectively, anti-CCRL2 ADC shows CCRL2-selective cytotoxicity against MDS/AML cell lines *in vitro* [21, 23, 24].

### Anti-CCRL2 ADC shows higher cytotoxicity against MDS/AML cell lines with erythroleukemic features and *TP53* mutations compared to gemtuzumab and PBC-conjugated ADCs targeting CD33 and CD123

Next, to compare the efficacy of ADC targeting CCRL2 with the efficacy of ADCs against other targets, PBD-conjugated ADCs targeting the well described AML targets CD33, and CD123 were developed (**Supplementary1E-F**). The cytotoxicity of the calichemicin-conjugated ADC gemtuzumab and the PBD-conjugated anti-CD33, -CD123, -CCRL2 ADCs were assessed in AML cell lines derived from patients with *de novo*/*TP53*-wild type AML (MOLM13, OCI-AML3 and MV4-11) (**Figure 2A-C**) and MDS/AML cell lines derived from patients with erythroleukemia, or MDS-related AML with *TP53* mutations (TF-1 - erythroleukemia, F36P – MDS-related erythroleukemia, and SKM1 – *TP53*-mutated MDS/AML) (**Figure 2D-F**). Consistently, with our previously published data, CCRL2 expression is significantly higher in MDS/AML cell lines with erythroleukemic features and *TP53* mutations compared to AML cell lines derived from patients with *de novo*/*TP53*-wild type AML (**Supplementary Figure 1G**).

**Figure 2.**
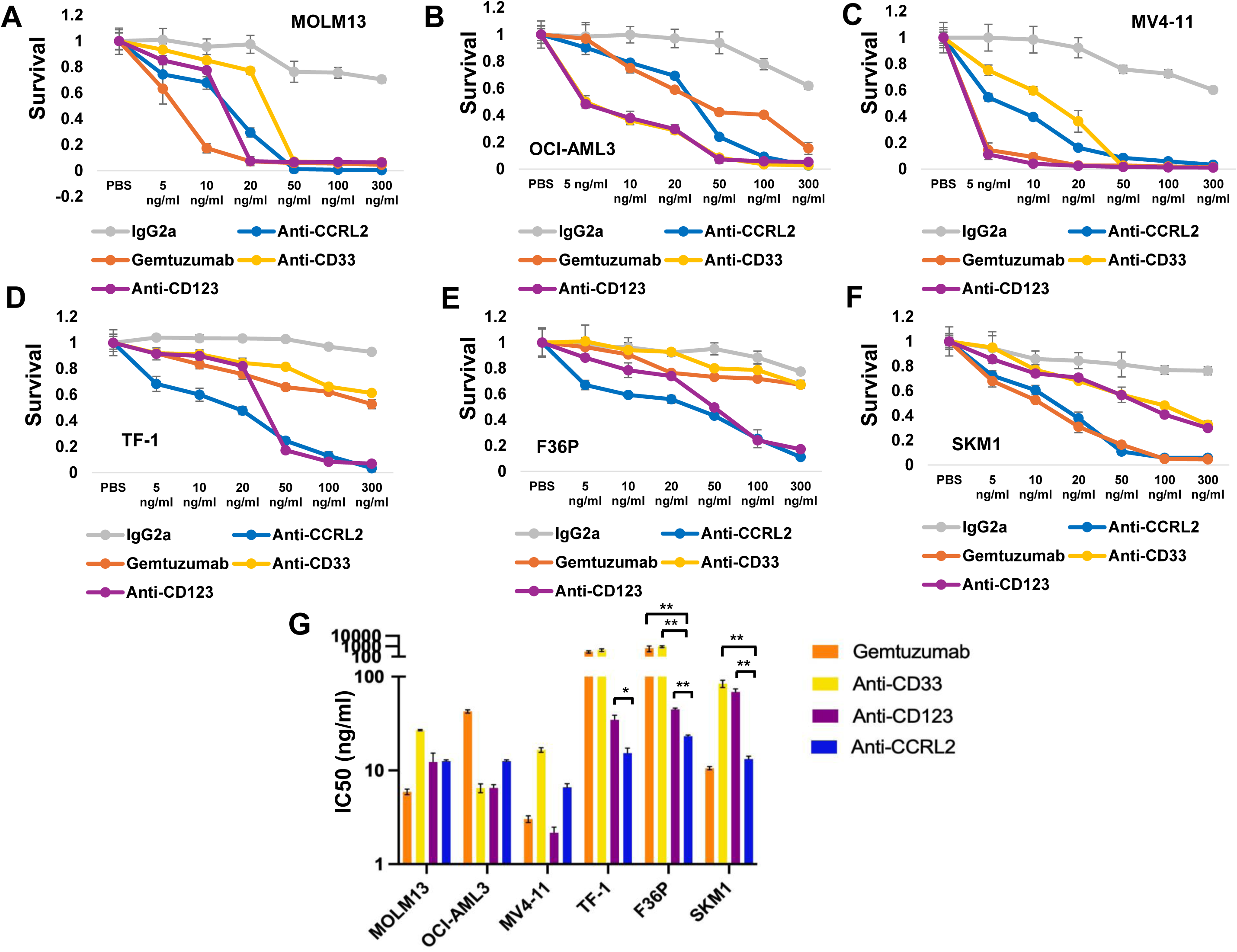
Anti-CCRL2 ADC shows higher cytotoxicity against MDS/AML cell lines with erythroleukemic features and *TP53* mutations compared to gemtuzumab and PBC-conjugated ADCs targeting CD33 and CD123. **(A-C)** Anti-CCRL2 ADC cytotoxicity against MOLM13, OCI-AML3 and MV4-11 cells derived from *de novo*/*TP53*-wild-type AML patients is relatively similar or lower compared to gemtuzumab and PBD-conjugated ADCs targeting CD33 and CD123. (**D**) Anti-CCRL2 ADC cytotoxicity against TF-1 cells (IC50 = 15.3 ng/ml) is higher compared to anti-CD123 (IC50 = 31.2 ng/ml), anti-CD33 (IC50 = 438.9 ng/ml) PBD-conjugated ADCs, gemtuzumab (IC50 = 296.5 ng/ml) or conjugated IgG2a (IC50 = 1159 ng/ml). (**E**) Anti-CCRL2 ADC cytotoxicity against F36P cells (IC50 =23.2 ng/ml) is higher compared to anti-CD123 (IC50 = 44.7 ng/ml), anti-CD33 (IC50 = 929.3 ng/ml) PBD-conjugated ADCs, gemtuzumab (IC50 = 1012.2 ng/ml) or conjugated IgG2a (IC50 = 1297.7 ng/ml). (**F**) Anti-CCRL2 ADC cytotoxicity against SKM1 cells (IC50 = 13.2 ng/ml) is higher compared to anti-CD123 (IC50 = 68.7 ng/ml), anti-CD33 (IC50 = 84.1 ng/ml) PBD-conjugated ADCs conjugated IgG2a (IC50 = 6455 ng/ml) and relatively lower compared to gemtuzumab (IC50 = 10.5 ng/ml). (G) The IC50 of anti-CCRL2 ADC is similar or higher compared to gemtuzumab and PBD-conjugated anti-CD33 and anti-CD123 ADCs in MOLM13, OCI-AML3 and MV4-11 cells but it is lower compared to gemtuzumab, PBD-conjugated anti-CD33 and anti-CD123 in TF-1 and F36P and compared to PBD-conjugated anti-CD33 and anti-CD123 in SKM1 cells. * p<0.010, ** p<0.001

AML cell lines derived from *de novo*/*TP53*-wild type AML were overall more sensitive to ADCs treatment compared to MDS/AML cell lines with erythroid features and *TP53* mutations (**Figure 2 A-C**). Particularly, the activity of gemtuzumab, and PBD-conjugated ADCs targeting CD33 and CD123 was found to be significantly decreased in MDS/AML cell lines with erythroid features and *TP53* mutations compared to AML cell lines derived from *de novo*/*TP53*-wild type AML. The anti-CCRL2 ADC showed overall similar or lower cytotoxicity compared to Gemtuzumab and PBD-conjugated ADCs targeting CD33 and CD123 in the AML cell lines derived from *de novo*/*TP53*-wild type AML (**Figure 2A, C**). However, within the MDS/AML cell lines with erythroid features and *TP53* mutations, the anti-CCRL2 ADC showed the highest cytotoxicity in TF-1 and F36P cells and the second to highest cytotoxicity in SKM1 cells (**Figure 2B, C**).

Overall, the activity of anti-CCRL2 ADC appears to be higher compared to other ADCs against MDS/AML cell lines with erythroid features and *TP53* mutations.

### Anti-CCRL2 ADC induces apoptosis and suppresses clonogenicity in MDS/AML primary cells

Next, primary mononuclear cells collected from 4 patients with *de novo/TP53*-wild-type AML, 4 patients with *TP53*-wild-type MDS-related AML and 10 patients with 10 multi-hit *TP53*-mutated (complex karyotype with *TP53* mutation with variant-allele frequency ≥50%) MDS/AML including 3 individuals with acute erythroid leukemia (**Supplementary Tables 2, 3 and Figure 3A**) were treated with 20 or 40 ng/ml anti-CCRL2 ADC or 20 ng/ml conjugated IgG2a for 5 days. Treatment with 10 and 20 ng/ml anti-CCRL2 increased apoptosis and suppressed cells clonogenicity compared to conjugated IgG2a (**Figure 3B**). The effect was relatively more prominent in *TP53*-mutated MDS/AML samples. Fluorescent in situ hybridization (FISH) of isolated colonies confirmed the presence of 17p and 5q deletions and trisomy 8 in isolated cells (**Supplementary Figure 2**). To assess if CCRL2 expression is associated with the cytotoxicity of the anti-CCRL2 ADC, the expression of CCRL2 in the surface of primary blasts was measured by flow cytometry (**Figure 3C**). Consistently with our previously reported results[14] [16], acute erythroid leukemia blasts express the highest levels of CCRL2 and TP53-mutated MDS/AML blasts express relatively higher levels of CCRL2 compared to *de novo* AML (**Figure 3C**). Linear regression showed a positive correlation between CCRL2 expression and anti-CCRL2 ADC apoptotic effect defined as the ratio of apoptosis induced by 40 ng/ml anti-CCRL2/apoptosis induced by 40 ng/ml IgG2a (**Figure 3D**).

**Figure 3.**
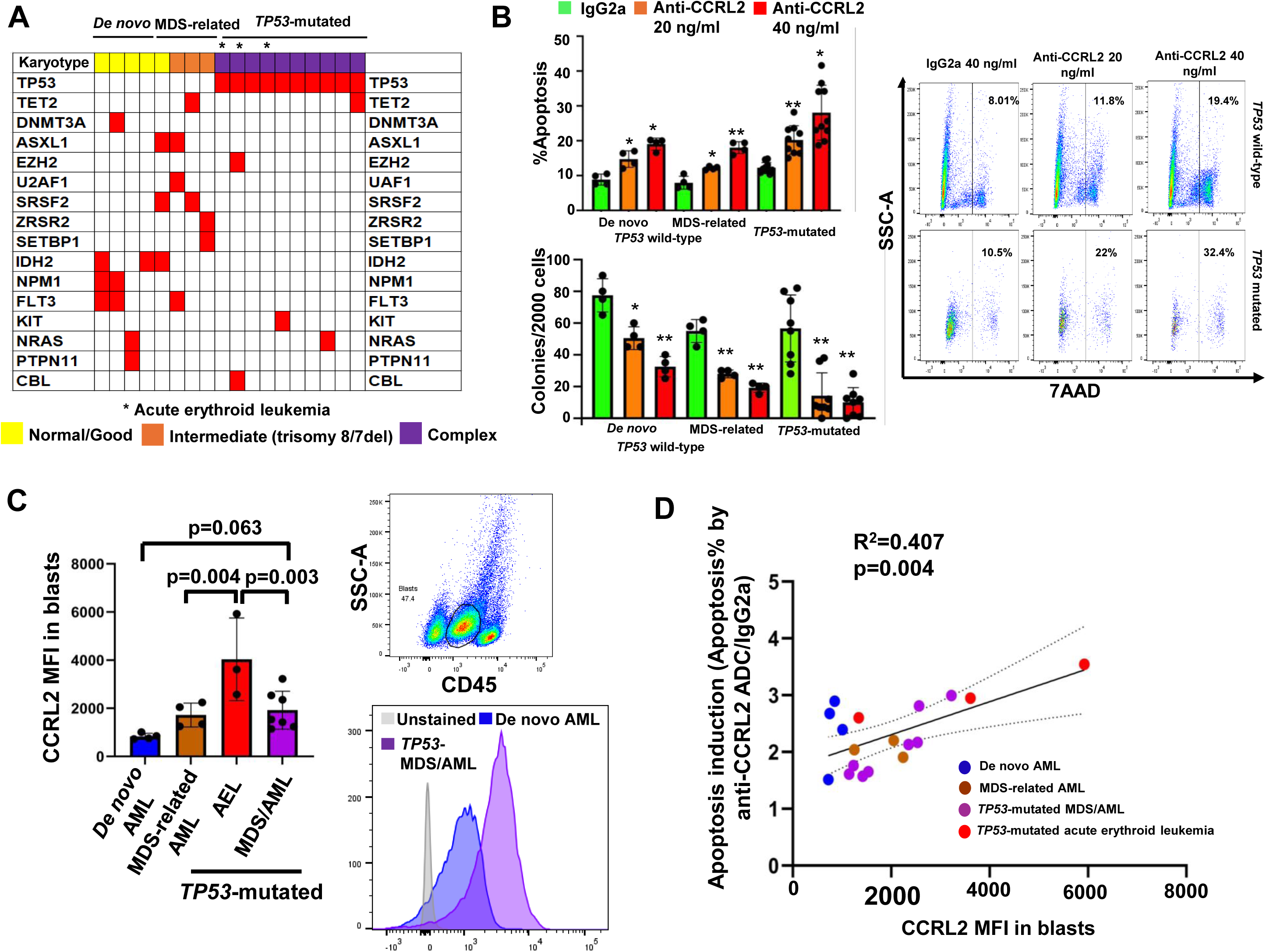
Anti-CCRL2 ADC induces apoptosis and suppresses clonogenicity in MDS/AML primary cells. (**A**) Genomic landscape of MDS/AML patients (*de novo TP53* wild-type, MDS-related *TP53* wild-type and *TP53*-mutated) included in the analysis of primary samples. (**B**) Treatment with 20 ng/ml and 40 ng/ml anti-CCRL2 ADC significantly increased apoptosis with impact being relatively more prominent in *TP53*-mutated samples. Treatment with 20 ng/ml and 40 ng/ml anti-CCRL2 ADC significantly suppressed clonogenicity of the primary MDS/AML samples with effect being overall more prominent in *TP53*-mutated samples. (**C**) CCRL2 expression in blasts gated by him CD45 and low SSC-A by flow cytometry is higher in *TP53* mutated acute erythroid leukemia patients compared to the rest of the groups and patients with *TP53* mutated MDS/AML have overall relatively higher CCRL2 expression compared to de novo AML (p=0.063). (**D**) The effect of anti-CCRL2 ADC in primary samples defined as the ratio of apoptosis with 40 ng/ml anti-CCRL2/apoptosis with 40 ng/ml IgG2a is positively associated with CCRL2 surface expression (R^2^=0.407, p=0.004). * p<0.01 and ** p<0.001.

Taken together, our results support that anti-CCRL2 ADC promotes apoptosis and suppresses the clonogenicity of primary MDS/AML cells. Higher CCRL2 expression, which is observed in *TP53*-mutated disease and particularly AML with erythroid features[16] is associated with better response.

### Anti-CCRL2 ADC does not affect healthy hematopoietic cells and has a good safety profile *in vivo*

To assess the effect of anti-CCRL2 ADC on healthy hematopoietic cells, CD34+ cells sorted from bone marrows of 3 independent healthy donors (**Supplementary Table 2**) were treated with 100 ng/ml conjugated IgG2a or anti-CCRL2 ADC, which is a substantially higher dose compared to the doses selected to treat primary MDS/AML samples for 5 days. Treatment with anti-CCRL2 ADC did not induce apoptosis (**Figure 4A**), affect differentiation (**Figure 4B**) or suppress clonogenicity (**Figure 4C**) of healthy CD34+ cells. Given that CCRL2 is overexpressed in leukemias with erythroid differentiation and its deletion decreases the survival and clonogenicity of erythroleukemic cells[16], the effect of anti-CCRL2 ADC on healthy erythroid progenitors was assessed. Mononuclear cells from 3 additional healthy bone marrow donors were treated with 100 ng/ml anti-CCRL2 or conjugated IgG2a for 5 days. Treatment with anti-CCRL2 ADC did not alter the percentages of erythroid progenitors CD71+CD235a-, CD71+CD235a+ or CD71-CD235a) (**Figure 4D**).

**Figure 4.**
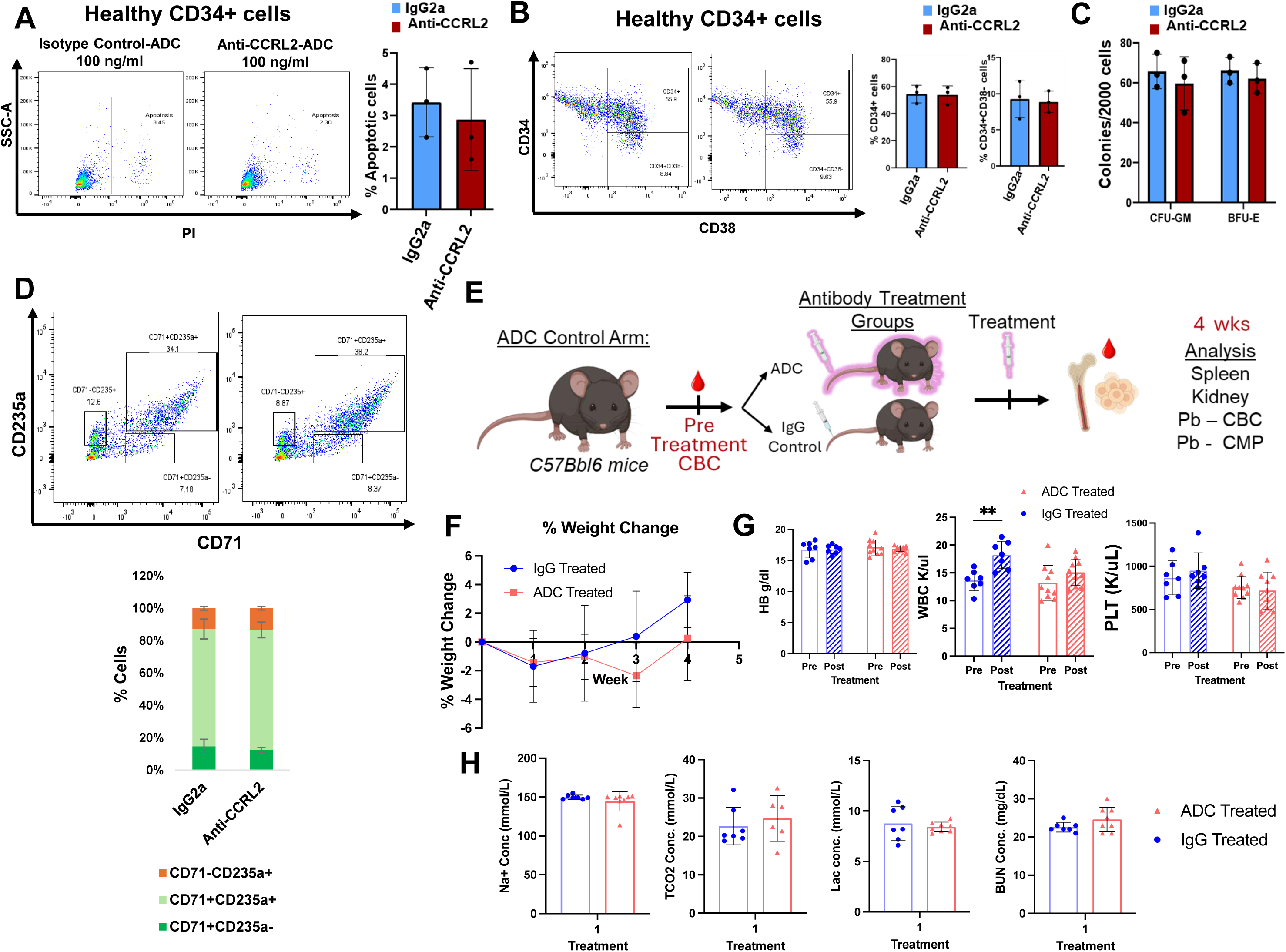
Anti-CCRL2 ADC does not affect healthy hematopoietic cells and has a good safety profile *in vivo*. Treatment of sorted CD34+ cells from two independent healthy bone marrow donors with 100 ng/ml of conjugated IgG2a or anti-CCRL2 ADC for 5 days did not increase apoptosis (**A**) and did not alter the CD34+% and CD34+CD38-% (**B**). **(C)** There was no impact on the clonogenicity of healthy CD34+ cells with treatment with 100 ng/ml anti-CCRL2 ADC. **(D)** Treatment of MNCs from 2 different healthy bone marrow donors with 100 ng/ml anti-CCRL2 ADC resulted in no significant alteration of the percentages of erythroid progenitors (CD71+CD235a-, CD71+CD235a+ or CD71-CD235a+). (**E**) Complete blood counts (CBC) were tested in C57BL/6 mice, which were then randomized to receive either anti-CCRL2 ADC or conjugated IgG2b (1 mg/kg, intravenously). Mice weight were measured weekly and CBC and chemistry panels were analyzed at 4 weeks after treatment. Mice were harvested and kidney and spleen weights were measured. (**F**) Weekly weight changes in mice treated with anti-CCRL2 ADC or conjugated IgG2b. (**G**) Pre- and post-Hemoglobin (Hgb), hematocrit (HCT), white blood cells (WBC) and platelets (PLT) are not different between anti-CCRL2 ADC or conjugated IgG2b treated mice. (**H**) Chemistry panels are not different between anti-CCRL2 ADC or conjugated IgG2b treated mice.

To further characterize the safety profile of this agent *in vivo*, an anti-mouse CCRL2 ADC was developed (**Supplementary Figure 3A**). Healthy 6-8 weeks C57BL/6 female mice were treated intravenously with 1 mg/kg anti-CCRL2 ADC or conjugated IgG2b (7 mice per group) (**Figure 4E**). No significant weight changes or systemic toxicities were observed following treatment (**Figure 4F**) and the weight of spleens and kidneys were not affected by treatment at 4 weeks following injection (**Supplementary Figure 3B**). Consistently, no significant alterations of the mice complete blood counts (**Figure 4G, Supplementary Figure 3C**) or chemistry panels (**Figure 4H, Supplementary Figure 3D**) were found.

Collectively, our results support that anti-CCRL2 ADC has a relatively good safety profile without prominent effect on healthy hematopoiesis.

### Anti-CCRL2 ADC suppresses leukemic growth and improves the survival in *TP53*-mutated MDS/AML cell line xenografts

To test the *in vivo* efficacy of one dose of anti-CCRL2 ADC in *TP53*-mutated MDS/AML models, TF-1 and SKM1 xenografts were treated. Luciferase+ TF-1 cells were intravenously injected in NSG mice. On day 15, to ensure balanced baseline tumor burdens, mice were stratified into groups with comparable bioluminescence signals and received anti-CCRL2 ADC or conjugated IgG2a (0.75 mg/kg, one dose) (**Figure 5A**). Mice treated with anti-CCRL2 ADC showed suppressed leukemic growth (**Figure 5B**) and improved survival compared to those treated with conjugated IgG2a (median survival 47 vs 37 days) (**Figure 5C**). Luciferase+ SKM1 cells were intravenously injected in NSG mice, which were randomized based on their bioluminescence signal on day 11 to anti-CCRL2 ADC or conjugated IgG2a (1 mg/kg, one dose) (**Figure 5D**). Treatment with anti-CCRL2 ADC suppressed leukemic growth (**Figure 5E**) and significantly prolonged the mice survival compared to those treated with conjugated IgG2a (median survival 39 vs 9.5 days) (**Figure 5F**). Of note, mice engrafted with SKM1 cells exhibit particularly enlarged livers at the time of their death. The mice treated with anti-CCRL2 ADC had significantly lower liver weights compared to those treated with IgG2a (**Figure 5G**).

**Figure 5.**
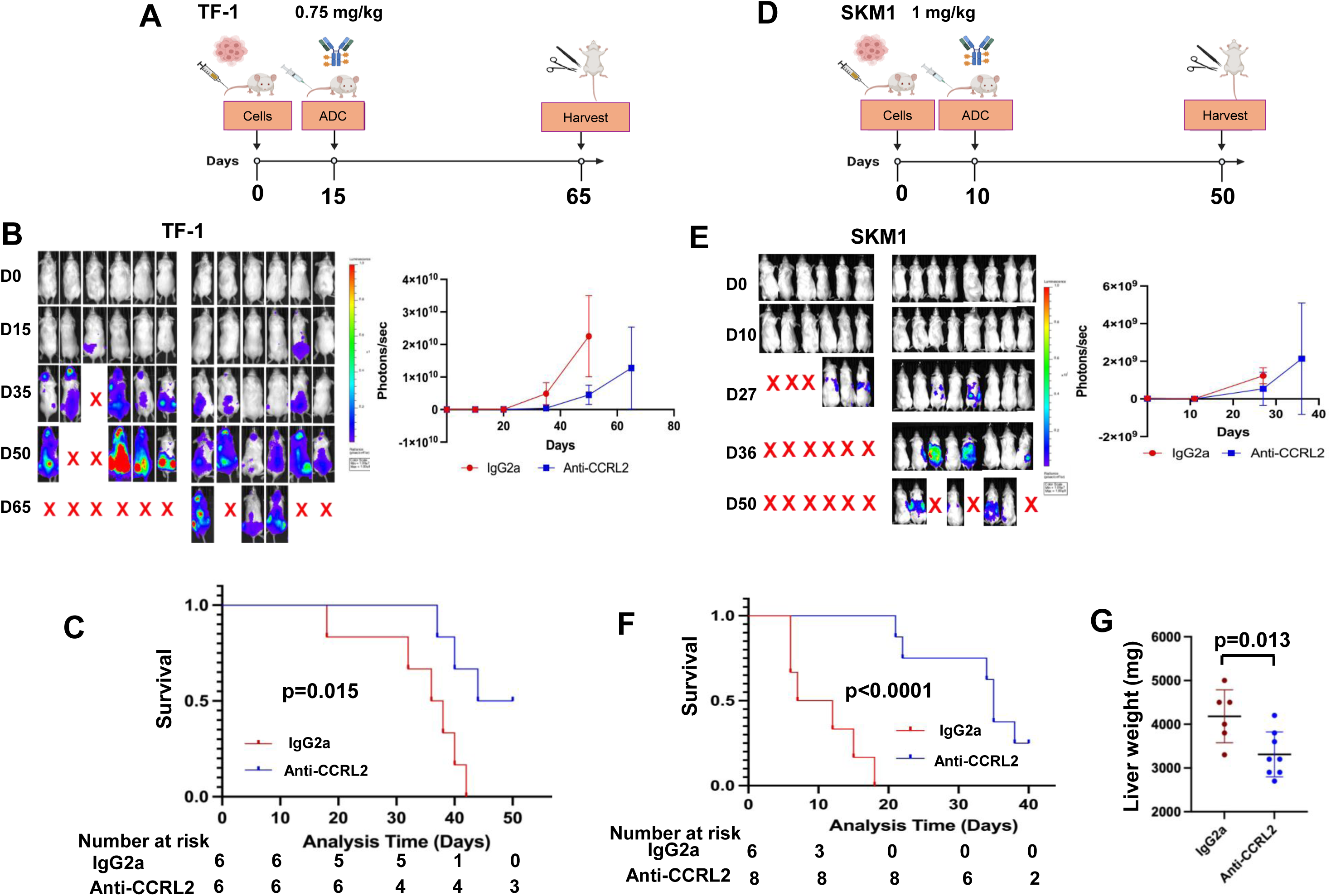
Anti-CCRL2 ADC suppresses the leukemic growth of TF-1 and SKM1 xenografts. **(A)** Luciferase+ TF-1 cells were injected intravenously in NSG mice (10^6^/mouse). At day 15 after injection, mice were randomized based on their bioluminescence signal to receive one intravenous dose of 0.75 mg/kg of anti-CCRL2 ADC or IgG2a ADC and monitored until day 65. **(B)** Mice treated with the anti-CCRL2 ADC showed suppressed leukemic growth by bioluminescence signal. (**C**) Mice treated with anti-CCRL2 ADC had improved overall survival (median survival 47 vs 37 days) compared to mice treated with IgG2a (p=0.015). (**D)** Luciferase+ SKM1 was injected intravenously in NSG mice (10^6^/mouse). At day 10 after injection, mice were randomized based on their bioluminescence signal to receive either one intravenous dose of 1 mg/kg of anti-CCRL2 ADC or IgG2a and monitored until day 50. (**E**) Mice treated with the anti-CCRL2 ADC showed decreased leukemic growth by bioluminescence signal compared to those treated with IgG2a. (**F**) Mice treated with anti-CCRL2 ADC showed significantly improved overall survival compared to mice treated with IgG2a (39 vs 9.5 days) (p<0.0001). (**G**) Mice treated with anti-CCRL2 ADC had lower liver weights compared to those treated with conjugated IgG2a.

Next, the effect of multiple anti-CCRL2 ADC doses was tested in the least aggressive *TP53*-mutated MDS/AML F36P xenograft model. At day 15 following intravenous injection of Luciferase+ F36P cells, mice received 1 mg/kg anti-CCRL2 ADC or conjugated IgG2a (**Figure 6A, Supplementary Figure 4A**). At day 35, mice were sacrificed, and those treated with anti-CCRL2 ADC had significantly lower percentage of human CD45+ cells (hCD45+%) in their bone marrows compared to those (**Figure 6B**). Another group of F36P xenografts received two doses of 1 mg/kg of anti-CCRL2 ADC or conjugated IgG2a at days 15 and 35 (**Figure 6C**). Mice treated with anti-CCRL2 ADC demonstrated suppressed leukemic growth (**Figure 6D**), improved survival (median survival 55 vs 38.5 days) (**Figure 6E**) and decreased hCD45+% in their bone marrows compared (**Figure 6F**) to those treated with conjugated IgG2a.

**Figure 6.**
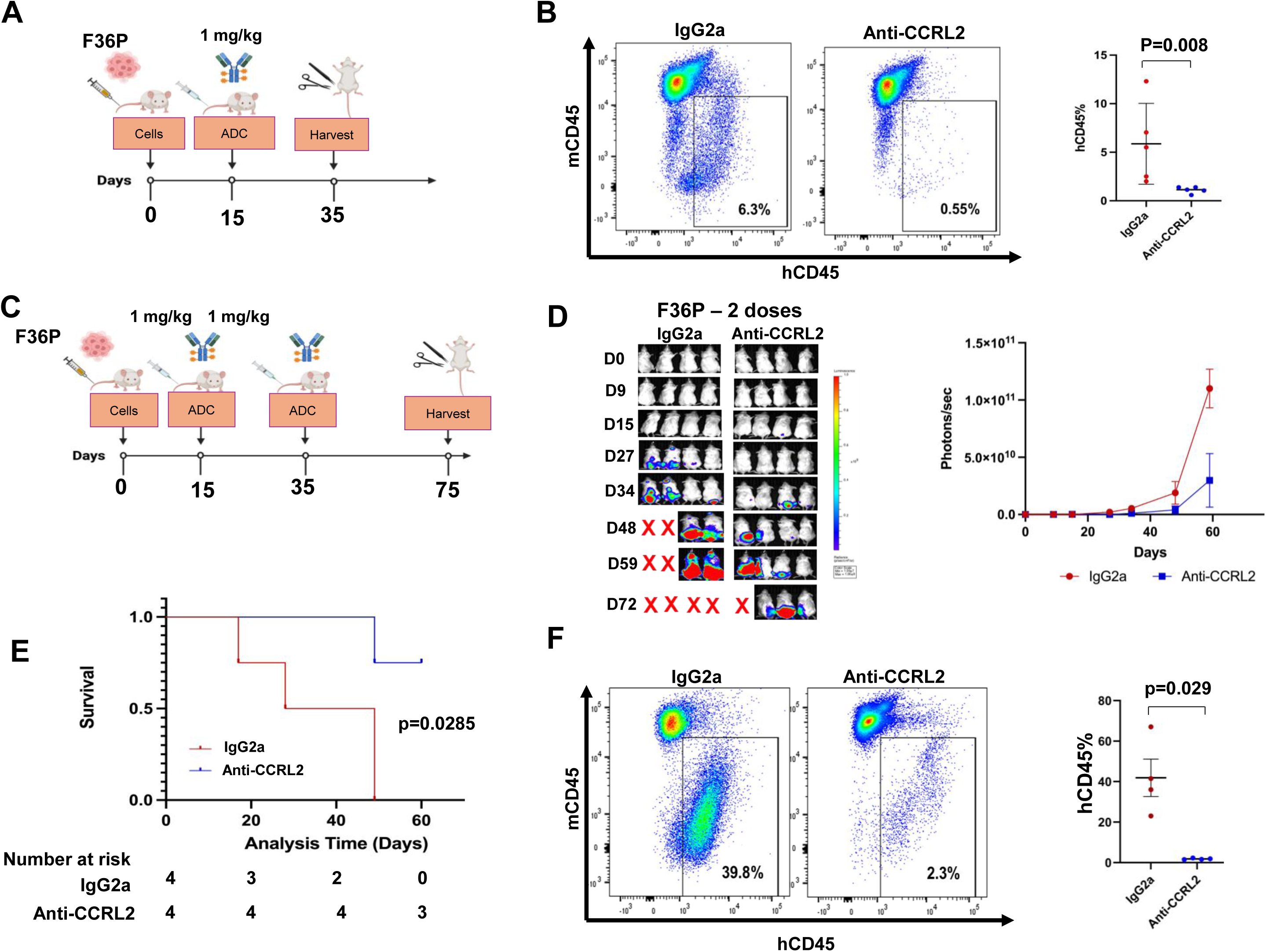
Multiple anti-CCRL2 ADC doses suppress the leukemic growth of F36P xenografts. **(A)** Luciferase+ F36P cells were injected intravenously in 10 NSG mice (10^6^/mouse) and at day 15, they were randomized to receive either one intravenous dose of 1 mg/kg of anti-CCRL2 ADC or IgG2a and were harvested at day 35. (**B**) Mice treated with anti-CCRL2 ADC had significantly lower percentage of human CD45+ cells in their bone marrow (p=0.008) compared to those treated with conjugated IgG2a. **(C)** Luciferase+ F36P cells were injected intravenously in 8 NSG mice (10^6^/mouse) and at day 15 they were randomized to receive either one intravenous dose of 1 mg/kg of anti-CCRL2 ADC or IgG2a. At day 35, mice received a second dose of 1 mg/kg of anti-CCRL2 ADC or IgG2a and were monitored until day 75. (**D**) Mice treated with the two anti-CCRL2 ADC doses showed suppressed leukemic growth by bioluminescence signal. (**E**) Mice treated with the two anti-CCRL2 ADC doses had significantly improved overall survival compared to mice treated with conjugated IgG2a (55 vs 38.5 days) (p=0.0285) with 3/4 mice surviving until day 75. Mice treated with the two anti-CCRL2 ADC doses had decreased percentage of human CD45+ cells in their bone marrows compared to conjugated IgG2a (p=0.021).

Taken together, these results support that treatment with anti-CCRL2 ADC suppresses the leukemic growth of *TP53*-mutated MDS/AML cell line xenografts.

### Anti-CCRL2 ADC suppresses the leukemic growth in multi-hit *TP53*-mutated MDS/AML patient-derived xenografts

Next, the efficacy of anti-CCRL2 ADC in patient-derived xenografts was assessed. Mononuclear cells from 5 *TP53-*mutated MDS/AML patients (**Supplementary Table 4**) were injected in sub-lethally irradiated (1.5 Gy) NSGS mice. Engraftment was assessed on day 30 by peripheral blood and bone marrow aspirate flow cytometry analysis (**Figure 7A**). Three out of the 5 patient samples were successfully engrafted (≥2% hCD45+ in bone marrow) in total 11 mice (**Figure 7A**). To ensure balanced baseline tumor burdens, mice were stratified into groups with comparable hCD45+% in their bone marrows, received anti-CCRL2 ADC or conjugated IgG2a (intravenously, 1 mg/kg) and harvested at day 55 (**Figure 7A**). The mice who were treated with anti-CCRL2 ADC showed a reduction or stabilization of hCD45+% in their peripheral blood (**Figure 7B**) or their bone marrow (**Figure 7C**) compared to those treated with conjugated IgG2a. Particularly, the bone marrow hCD45+% fold change was significantly lower in mice treated with anti-CCRL2 ADC compared to those treated with conjugated IgG2a (**Figure 7C**). Similarly, mice treated with anti-CCRL2 ADC had significantly lower hCD45+% in their spleens compared to those treated with conjugated IgG2a (**Figure 7D**).

**Figure 7.**
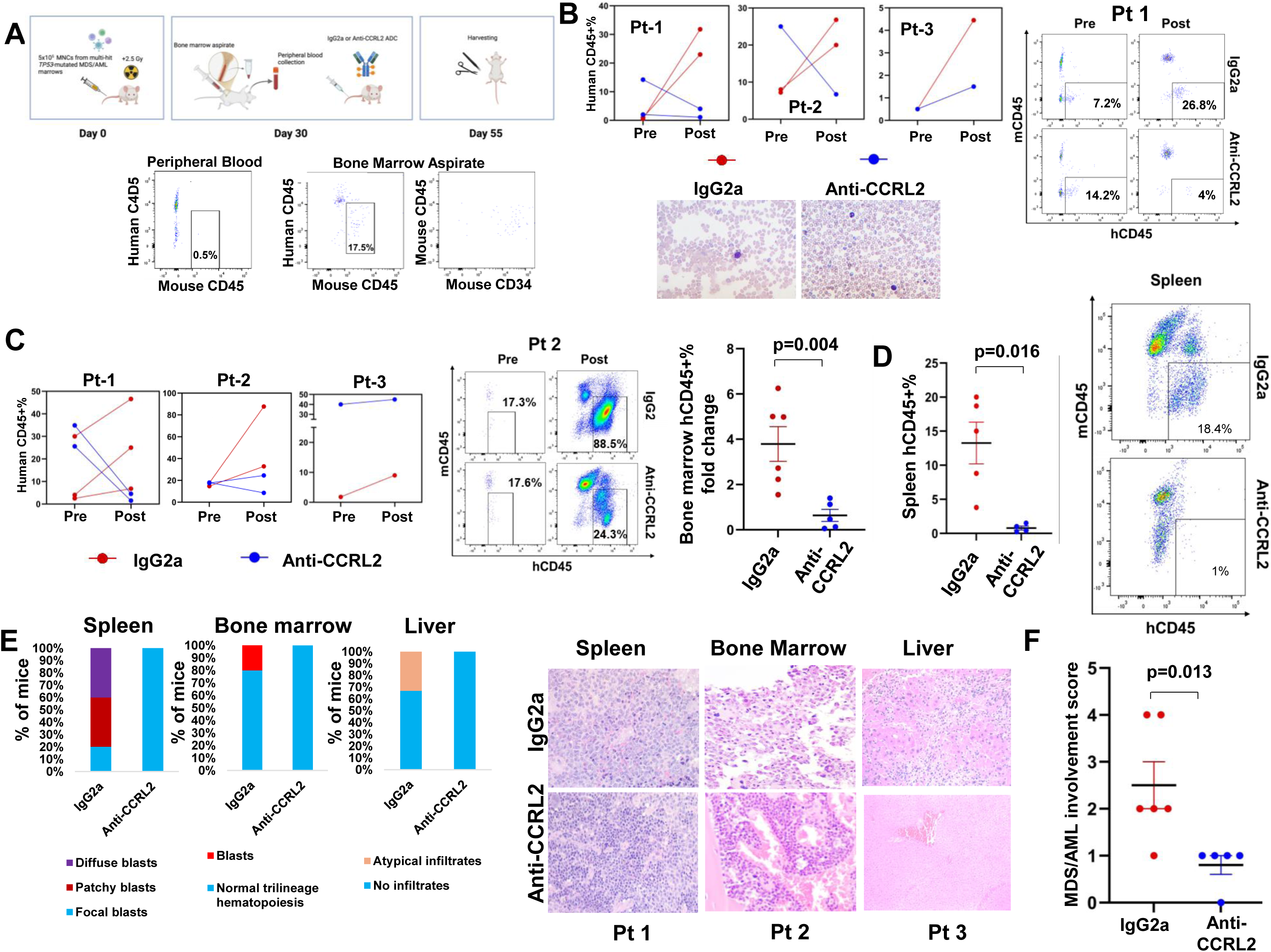
Anti-CCRL2 ADC suppresses the leukemic growth in multi-hit *TP53*-mutated MDS/AML patient-derived xenografts. **(A)** A scheme summarizing the development and treatment of multi-hit *TP53*-mutated MDS/AML patient-derived xenografts. MNCs isolated from bone marrow aspirates of patients with *TP53*-mutated MDS/AML containing leukemic blasts were injected intravenously in NSGS mice (5×10^5^/mouse). At day 30, peripheral blood and bone marrow aspirates were collected from mice and engraftment was assessed (≥2% of human CD45+ cells in the bone marrow). Human cells were almost uniformly CD34+. Mice were then randomized based on their human CD45+% to receive intravenously either one dose of 1 mg/kg anti-CCRL2 ADC or conjugated IgG2a. Twenty-five days later mice were sacrificed to assess disease burden. **(B)** Mice treated with the anti-CCRL2 ADC showed a decline or stabilization of human CD45+% in blood as opposed to mice treated with conjugated IgG2a, which showed a prominent increase in human CD45+%. Representative peripheral blood smear showing circulating blasts in an IgG2a treated mouse, in parallel to differentiated leukocytes in an anti-CCRL2 ADC treated mouse. **(C)** Mice treated with the anti-CCRL2 ADC showed a decline or stabilization of bone marrow human CD45+% as opposed to mice treated with conjugated IgG2a, which showed a prominent increase in bone marrow human CD45+% cells before and after treatment. Mice treated with anti-CCRL2 ADC showed a significantly lower fold increase of human CD45+% in their bone marrows compared to those treated with IgG2a (p=0.004). (**D**) Mice treated with anti-CCRL2 ADC had a relatively lower percentage of human CD45+ cells in their spleens compared to those treated with conjugated IgG2a (p=0.016). (**E**) Immunohistochemical analysis of spleens, bone marrows and livers showed lower leukemic involvement in mice treated with anti-CCRL2 ADC compared to IgG2a. Representative figures showing diffuse involvement of leukemic blasts (IgG2a) as opposed to patchy involvement (anti-CCRL2 ADC) (mice engrafted with patient 1 MNCs), leukemic blasts (IgG2a) as opposed to normal bone marrow (anti-CCRL2 ADC) (mice engrafted with patient 2 MNCs) and atypical infiltrates in the liver (IgG2a) as opposed to normal liver (anti-CCRL2 ADC) (mice engrafted with patient 3 MNCs). (**F**) Mice treated with anti-CCRL2 ADC had a significantly lower MDS/AML involvement score (p=0.013) compared to those treated with conjugated IgG2a.

Pathologic analysis of mouse tissues revealed that mice treated with conjugated IgG2a had more advanced disease involvement in their marrows, spleens and livers compared to those treated with anti-CCRL2 ADC (**Figure 7E**). An MDS/AML involvement score was developed (**Supplementary Figure 4B**) and mice treated with anti-CCRL2 ADC had significantly lower scores compared to those treated with conjugated IgG2a (**Figure 7F**).

Collectively, these results suggest that treatment with anti-CCRL2 ADC suppresses leukemic growth in multi-hit *TP53*-mutated patient-derived MDS/AML xenografts.

## Discussion

Patients with high-risk MDS/AML including those with *TP53* mutations and deletions continue to exhibit relatively poor outcomes due to the lack of effective therapies and high incidence of treatment resistance[1–3]. ADCs including the PBD-conjugated loncastuximab, have prominent activity in hematologic malignancies [8, 9, 21, 22], but the currently available ADCs such as the anti-CD33 ADC gemtuzumab, show limited activity in high-risk MDS/AML and *TP53*-mutated disease [10]. We developed an anti-CCRL2 ADC using the PBD payload and showed a significant single-agent anti-leukemic effect in various MDS/AML models.

Our studies demonstrated that cell lines derived from patients with high-risk MDS/AML or AML with erythroid features are generally less sensitive to gemtuzumab and PBD-conjugated ADCs targeting CD33 and CD123, compared to AML cell lines from patients with *de novo* or good-to intermediate-risk disease. This finding aligns with the clinical responses observed in high-risk MDS/AML patients treated with gemtuzumab[10]. In contrast, the anti-CCRL2 ADC exhibited consistent efficacy across the different cell line subtypes, demonstrating higher activity against cell lines derived from patients with *TP53* mutated MDS/AML or AML with erythroid features compared to gemtuzumab and PBD-conjugated ADCs targeting CD33 and CD123. These results suggest that the marked upregulation of CCRL2 in high-risk MDS/AML and *TP53*-mutated cells may offset their inherent resistance to drug toxin-mediated cell killing.

Analysis of anti-CCRL2 ADC activity in primary samples confirmed that this agent exerts a prominent pro-apoptotic effect and significantly suppresses the clonogenic potential of MDS/AML cells, with the highest efficacy observed in *TP53*-mutated MDS/AML samples and particularly acute erythroid leukemia samples. Notably, there was a positive association between CCRL2 expression and the apoptotic activity induced by the agent, consistent with findings reported for other ADCs targeting CD33[25–27]. Our group recently demonstrated that *TP53*-mutated MDS/AML and AML with erythroid features exhibit the highest levels of CCRL2 expression across the spectrum of MDS/AML subtypes[16]. These results may underlie the significant anti-leukemic effect of anti-CCRL2 ADC observed in erythroleukemic cell lines and in primary samples from patients with acute erythroid leukemia included in our analysis.

PBD-conjugated antibody-drug conjugates (ADCs) targeting CD33 and CD123 showed promising preclinical efficacy in high-risk AML[28, 29]. However, early-phase clinical trials revealed cytopenias and hepatotoxicity, which hindered further clinical development. Notably, not only gemtuzumab but also bi-specific antibodies and CAR-T cells targeting CD33, have been associated with cytopenias[30–32]. This suggests that these adverse effects may be related to the expression of CD33 and CD123 on the surface of healthy hematopoietic cells. While transient elevations in liver enzymes have been observed with PBD payloads, there have been no reports of fatal hepatotoxicity[21, 22]. On the contrary, healthy stem and progenitor cells express minimal CCRL2 levels[14, 15]. Consistently, anti-CCRL2 ADCs demonstrated minimal impact on healthy hematopoietic cells, and did not induce systemic toxicity or significant changes in blood counts or chemistry panels in C57BL/6 mice, similarly to recent findings from our group[33]. Should cytopenias or hepatotoxicity arise with anti-CCRL2 ADCs, fractionated dosing may be considered, as this approach has enabled the safe and effective use of gemtuzumab.

Our *in vivo* studies demonstrated that the anti-CCRL2 ADC effectively suppressed leukemic growth in cell line xenograft models, resulting in improved survival of treated mice. Notably, two doses of the agent administered 20 days apart were well tolerated by NSG mice and led to a marked reduction in disease burden. Furthermore, bone marrow aspirates collected from NSGS mice engrafted with *TP53*-mutated MDS/AML cells enabled randomization to treatment with either anti-CCRL2 ADC or an isotype control. This approach revealed significant anti-leukemic activity of the anti-CCRL2 ADC.

Further studies in syngeneic MDS/AML models are required to solidify the efficacy and safety of anti-CCRL2 ADC, but this agent appears to be a promising candidate for single-agent or combination therapies particularly for high-risk MDS/AML including *TP53*-mutated disease. Similarly, conjugating antibodies targeting other surface antigens that are particularly overexpressed in high-risk MDS/AML and show very low expression in healthy hematopoietic cells, with PBD may be an attractive strategy to improve the outcomes of patients with these neoplasms.

## Author contributions

N.S.N., S.P. and T.K. conceived and designed the study. N.S.N. and T.K. wrote the manuscript. T.A., B.S.L., X.Z, T.D.N. and S.P. performed the antibody-drug conjugation. N.S.N., B.P., T.C. Y.A., X.Z. and T.K. performed the MTT assays, apoptotic assays and clonogenicity assays with cell lines. B.P. and T.K. processed primary samples and performed the primary cell *in vitro* studies. M.K. and B.P. performed the FISH analysis, Z.T., L.L. and L.R. performed the safety studies in C57BL/6 mice. N.S.N, E.W., S.K., S.P. and T.K. performed the cell-line xenograft studies. N.S.N, E.W. G.G., T.S., S.K. and T.K. performed the patient-derived xenograft studies. D.J.P. performed the immunohistochemical analysis. I.G., A.E.D., M.J.L., and R.J.J. identified clinical samples and provided clinical information. S.P., T.K., A.E.D., I.G., G.G., M.J.L., R.J.J., S.K. interpreted the data and edited the manuscript.

## References

1. Kröger, N., Treatment of high-risk myelodysplastic syndromes. Haematologica, 2025. 110(2): p. 339–349.

2. Daver, N.G., et al., TP53-Mutated Myelodysplastic Syndrome and Acute Myeloid Leukemia: Biology, Current Therapy, and Future Directions. Cancer Discov, 2022. 12(11): p. 2516–2529.

3. Hochman, M.J., et al., Prognostic impact of secondary versus de novo ontogeny in acute myeloid leukemia is accounted for by the European LeukemiaNet 2022 risk classification. Leukemia, 2023. 37(9): p. 1915–1918.

4. Fu, Z., et al., Antibody drug conjugate: the "biological missile" for targeted cancer therapy. Signal Transduct Target Ther, 2022. 7(1): p. 93.

5. Paul, S., et al., Cancer therapy with antibodies. Nat Rev Cancer, 2024. 24(6): p. 399–426.

6. Calabretta, E., et al., The antibody-drug conjugate loncastuximab tesirine for the treatment of diffuse large B-cell lymphoma. Blood, 2022. 140(4): p. 303–308.

7. Kantarjian, H.M., et al., Inotuzumab Ozogamicin versus Standard Therapy for Acute Lymphoblastic Leukemia. N Engl J Med, 2016. 375(8): p. 740–53.

8. Freeman, S.D., et al., Fractionated vs single-dose gemtuzumab ozogamicin with determinants of benefit in older patients with AML: the UK NCRI AML18 trial. Blood, 2023. 142(20): p. 1697–1707.

9. Kovtun, Y., et al., A CD123-targeting antibody-drug conjugate, IMGN632, designed to eradicate AML while sparing normal bone marrow cells. Blood Adv, 2018. 2(8): p. 848–858.

10. Awada, H., et al., Gemtuzumab ozogamicin plus standard induction hemotherapy improves outcomes of newly diagnosed intermediate cytogenetic risk acute myeloid leukemia. Blood Cancer J, 2023. 13(1): p. 131.

11. Schioppa, T., et al., Molecular Basis for CCRL2 Regulation of Leukocyte Migration. Front Cell Dev Biol, 2020. 8: p. 615031.

12. Del Prete, A., et al., The Atypical Receptor CCRL2 Is Essential for Lung Cancer Immune Surveillance. Cancer Immunol Res, 2019. 7(11): p. 1775–1788.

13. Del Prete, A., et al., The atypical receptor CCRL2 is required for CXCR2-dependent neutrophil recruitment and tissue damage. Blood, 2017. 130(10): p. 1223–1234.

14. Karantanos, T., et al., The role of the atypical chemokine receptor CCRL2 in myelodysplastic syndrome and secondary acute myeloid leukemia. Sci Adv, 2022. 8(7): p. eabl8952.

15. Karantanos, T., et al., CCRL2 affects the sensitivity of myelodysplastic syndrome and secondary acute myeloid leukemia cells to azacitidine. Haematologica, 2023. 108(7): p. 1886–1899.

16. Naji, N.S., et al., *CCRL2 promotes the interferon-*γ *signaling response in myeloid neoplasms with erythroid differentiation and mutated TP53*. bioRxiv, 2025.

17. Rodriguez-Meira, A., et al., Single-cell multi-omics identifies chronic inflammation as a driver of TP53-mutant leukemic evolution. Nat Genet, 2023. 55(9): p. 1531–1541.

18. Wang, B., et al., *Comprehensive characterization of IFN*γ *signaling in acute myeloid leukemia reveals prognostic and therapeutic strategies*. Nat Commun, 2024. 15(1): p. 1821.

19. Nichakawade, T.D., et al., TRBC1-targeting antibody-drug conjugates for the treatment of T cell cancers. Nature, 2024. 628(8007): p. 416–423.

20. Hartley, J.A., et al., Pre-clinical pharmacology and mechanism of action of SG3199, the pyrrolobenzodiazepine (PBD) dimer warhead component of antibody-drug conjugate (ADC) payload tesirine. Scientific Reports, 2018. 8(1): p. 10479.

21. Zammarchi, F., et al., ADCT-402, a PBD dimer-containing antibody drug conjugate targeting CD19-expressing malignancies. Blood, 2018. 131(10): p. 1094–1105.

22. Hamadani, M., et al., Final results of a phase 1 study of loncastuximab tesirine in relapsed/refractory B-cell non-Hodgkin lymphoma. Blood, 2021. 137(19): p. 2634–2645.

23. Ogitani, Y., et al., DS-8201a, A Novel HER2-Targeting ADC with a Novel DNA Topoisomerase I Inhibitor, Demonstrates a Promising Antitumor Efficacy with Differentiation from T-DM1. Clin Cancer Res, 2016. 22(20): p. 5097–5108.

24. Tashakori, M., et al., Differential characteristics of TP53 alterations in pure erythroid leukemia arising after exposure to cytotoxic therapy. Leuk Res, 2022. 118: p. 106860.

25. Khan, N., et al., Expression of CD33 is a predictive factor for effect of gemtuzumab ozogamicin at different doses in adult acute myeloid leukaemia. Leukemia, 2017. 31(5): p. 1059–1068.

26. Olombel, G., et al., The level of blast CD33 expression positively impacts the effect of gemtuzumab ozogamicin in patients with acute myeloid leukemia. Blood, 2016. 127(17): p. 2157–60.

27. Kantarjian, H.M., et al., Inotuzumab Ozogamicin for Relapsed/Refractory Acute Lymphoblastic Leukemia in the INO-VATE Trial: CD22 Pharmacodynamics, Efficacy, and Safety by Baseline CD22. Clin Cancer Res, 2021. 27(10): p. 2742–2754.

28. Stein, E.M., et al., A phase 1 trial of vadastuximab talirine as monotherapy in patients with CD33-positive acute myeloid leukemia. Blood, 2018. 131(4): p. 387–396.

29. Kung Sutherland, M.S., et al., SGN-CD33A: a novel CD33-targeting antibody-drug conjugate using a pyrrolobenzodiazepine dimer is active in models of drug-resistant AML. Blood, 2013. 122(8): p. 1455–63.

30. Castaigne, S., et al., Effect of gemtuzumab ozogamicin on survival of adult patients with de-novo acute myeloid leukaemia (ALFA-0701): a randomised, open-label, phase 3 study. Lancet, 2012. 379(9825): p. 1508–16.

31. Ravandi, F., et al., Updated results from phase I dose-escalation study of AMG 330, a bispecific T-cell engager molecule, in patients with relapsed/refractory acute myeloid leukemia (R/R AML). Journal of Clinical Oncology, 2020. 38: p. 7508–7508.

32. Shah, N.N., et al., CD33 CAR T-Cells (CD33CART) for Children and Young Adults with Relapsed/Refractory AML: Dose-Escalation Results from a Phase I/II Multicenter Trial. Blood, 2023. 142(Supplement 1): p. 771–771.

33. Wang, T., et al., Targeting CCRL2 enhances therapeutic outcomes in a tuberculosis mouse model. Front Immunol, 2025. 16: p. 1501329.

