## Supplementary Figures for "A PBD-dimer containing antibody drug conjugate targeting CCRL2 for high-risk MDS/AML"

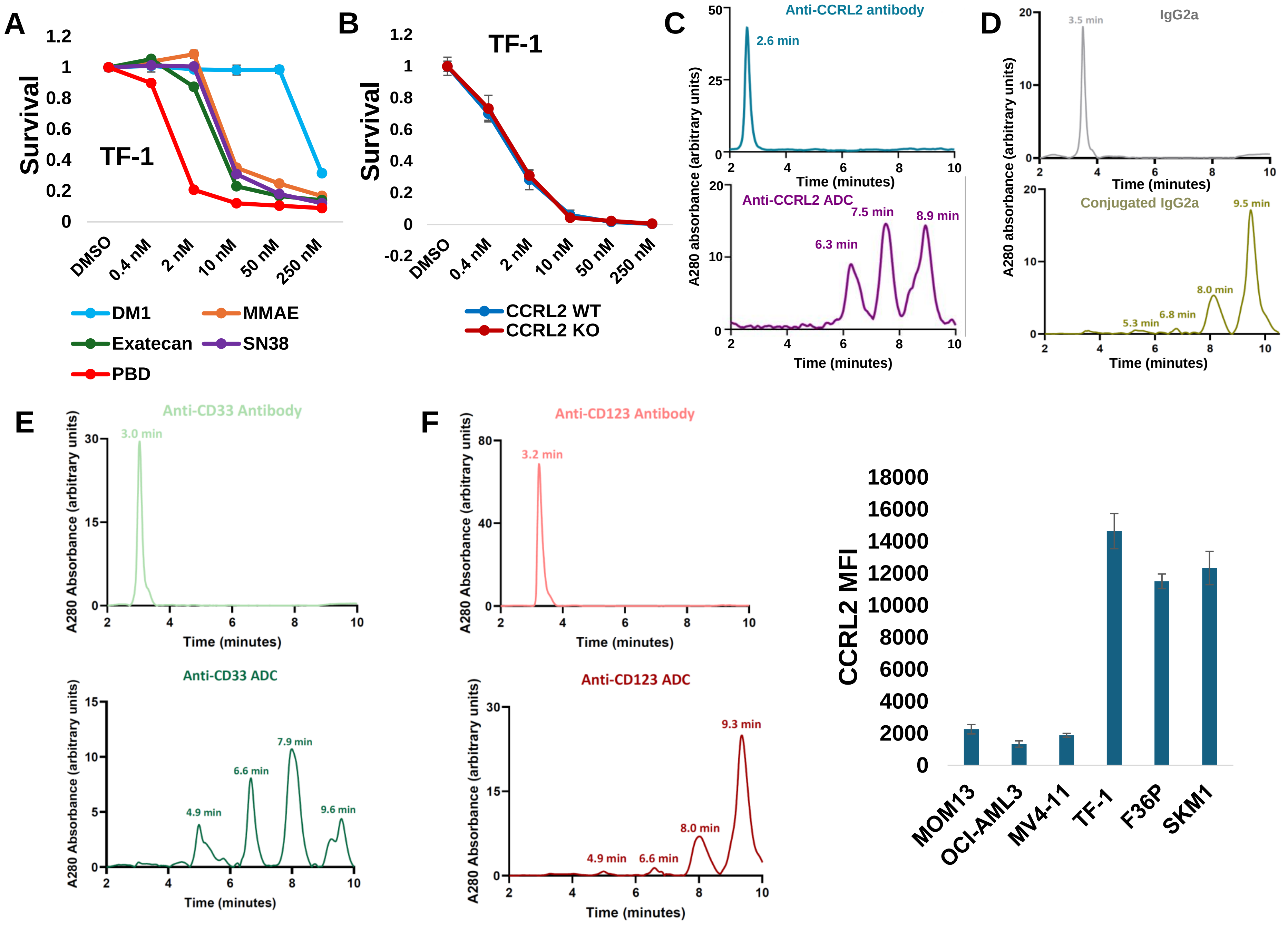


Supplementary Figure 1

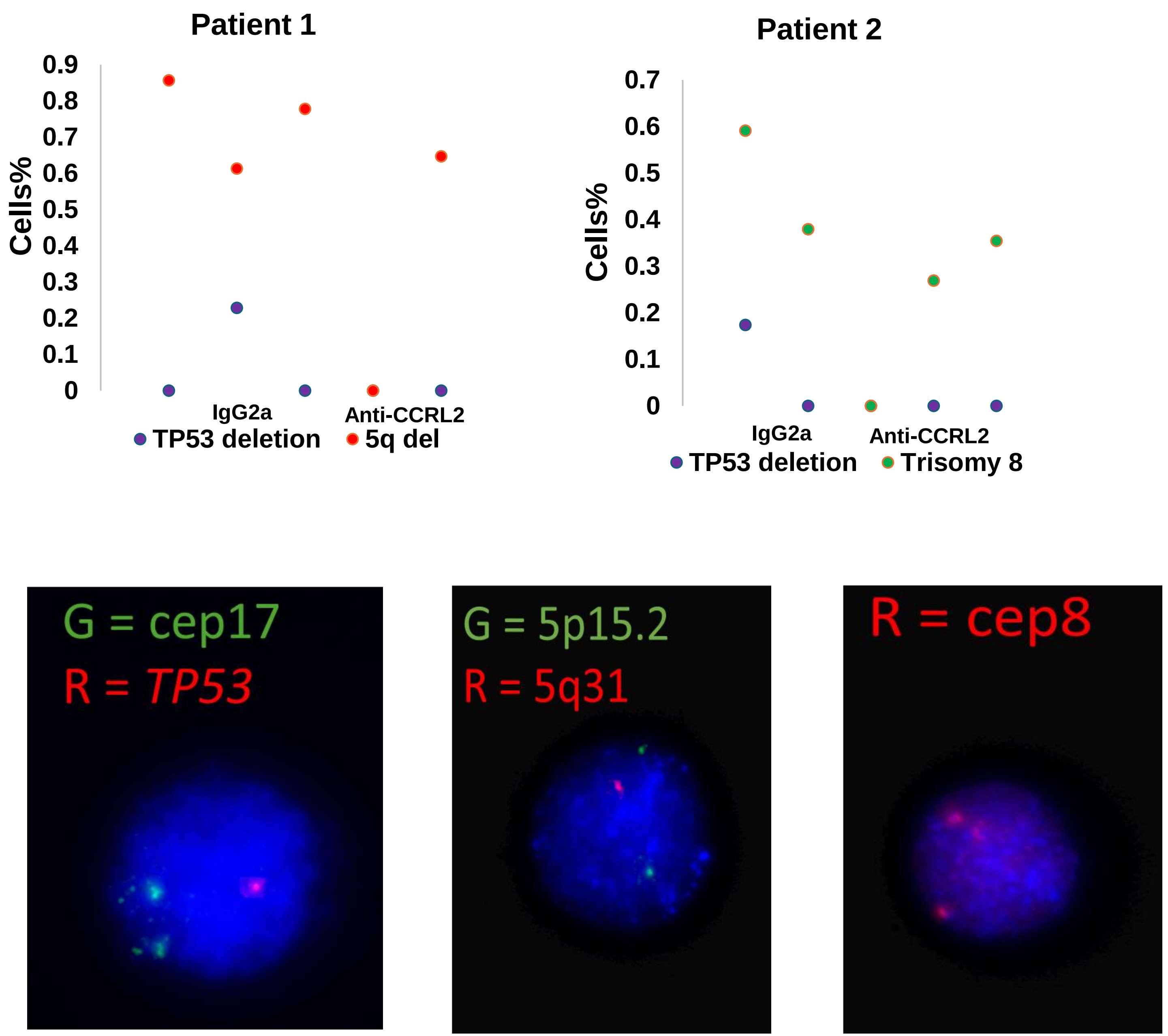


Supplementary Figure 2

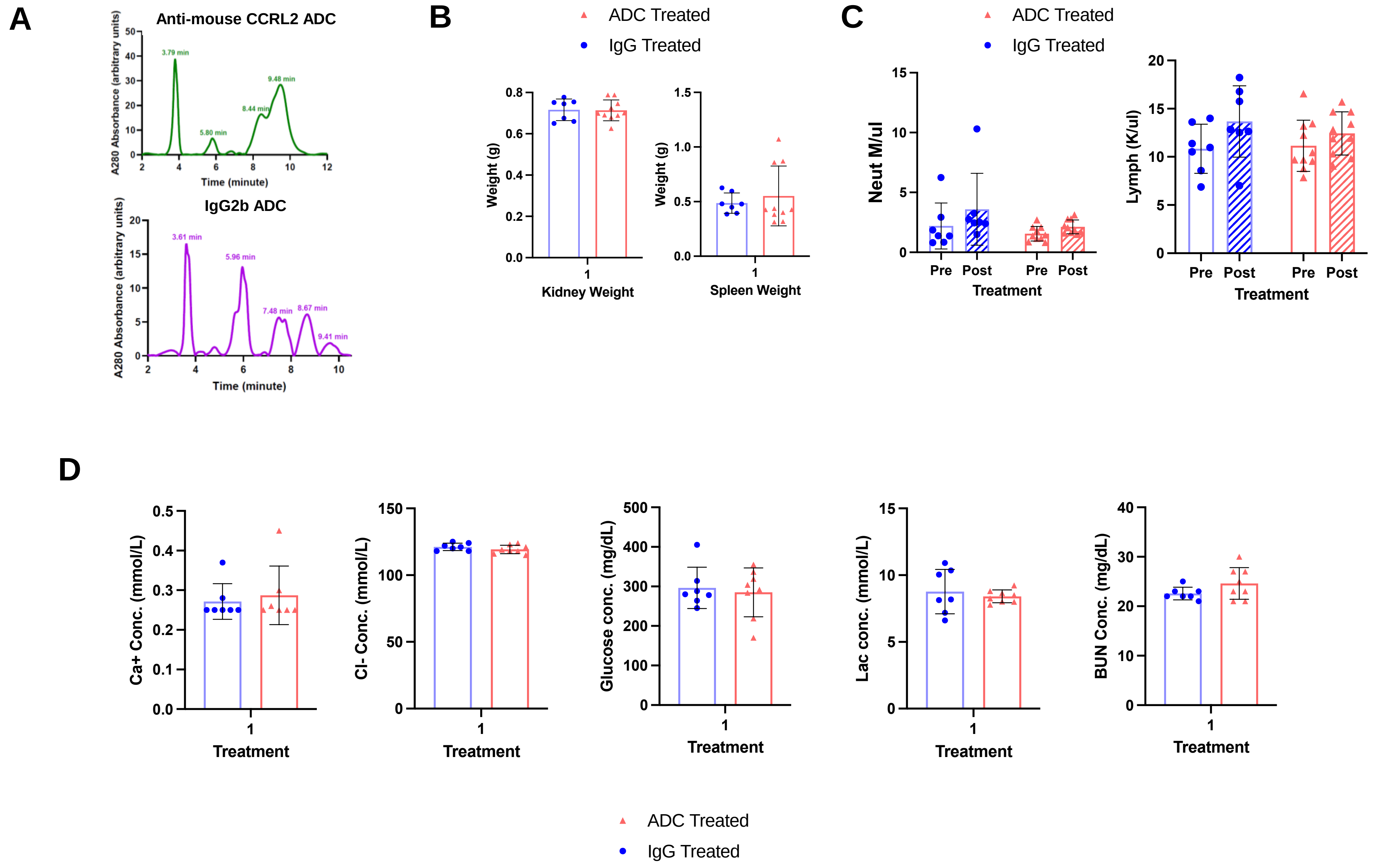


**Supplementary Figure 3**

A

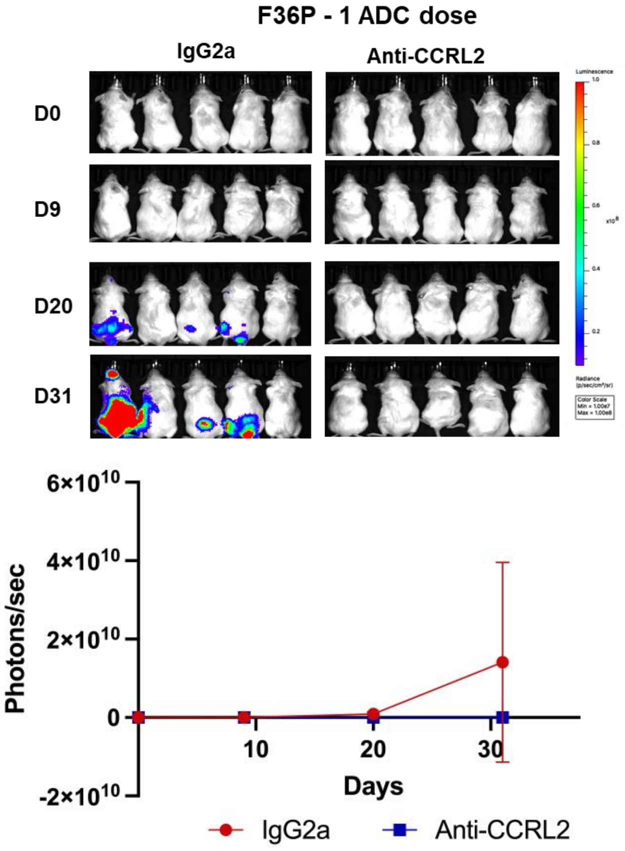


B

### MDS/AML involvement score

| Bone Marrow | Spleen | Liver |
|---|---|---|
| Normal 0 point | No blasts 0 point | Normal 0 point |
| Dysplasia 1 point | Focal blasts 1 | Atypical infiltrates 1 point |
| Blasts 1 point | Patchy blasts 2 | |
| | Diffuse/sheets 3 | |

**Supplementary Figure 4**
