## Supplementary Figure Legends for "A PBD-dimer containing antibody drug conjugate targeting CCRL2 for high-risk MDS/AML"

**Supplementary Figure 1. (A)** SG3199 showed significantly higher cytotoxicity against TF-1 cells compared to other drug toxins commonly used for ADC production (DM1, MME, Exatecan, SN38). (**B**) CCRL2 KO does not affect the sensitivity of TF-1 cell to SG3199. Development of ADCs by conjugating anti-CCRL2 (**C**), IgG2a (**D**), anti-CD33 (**E**) and anti-CD123 (**F**). The antibody-drug conjugation was confirmed by high-performance liquid chromatography. (**G**) Flow cytometry analysis shows that cell lines derived from patients with *TP53*-mutated MDS/AML or erythroleukemia (TF-1, F36P and SKM1) express significantly higher surface CCRL2 compared to cell lines derived from patients with *de novo*/*TP53*-wild type AML (MOLM13, OCI-AML3 and MV4-11) cells.

**Supplementary Figure 2.** Single colonies from primary cells from two independent *TP53*-mutated MDS/AML with complex karyotype treated with IgG2a (N=3 from patient 1 and 2 from patient 2) or anti-CCRL2 ADC (N=2 from patient 1 and 3 from patient 2) were analyzed by fluorescence in situ hybridization (FISH) identifying chromosomal abnormalities (*TP53* deletion, 5q deletion and trisomy 8) reported in the clinical samples at a significant percentage of tested cells.

**Supplementary Figure 3.** (**A**) Development of the anti-mouse CCRL2 ADC by conjugating anti-mouse CCRL2 antibody or IgG2b with pyrrolobenzodiazepine (PBD). (**B**) Kidney and spleen weights were not different between anti-CCRL2 ADC or IgG2b treated mice. (**C**) Absolute neutrophils and lymphocytes were not significantly different between mice treated with anti-CCRL2 ADC or conjugated IgG2b. (**D**) Chemistry panels are not different between anti-CCRL2 ADC or conjugated IgG2b treated mice.

**Supplementary Figure 4. (A)** Bioluminescence signal of NSG mice engrafted with F36P cells, treated with 1 mg/kg anti-CCRL2 ADC or IgG2a ADC at day 15 and harvested at day 35 to assess disease burden. (**B**) MDS/AML involvement score developed based on involvement of spleen, bone marrow and liver by leukemic cells.
