## Supplementary Tables for "A PBD-dimer containing antibody drug conjugate targeting CCRL2 for high-risk MDS/AML"

**Supplementary Table 1. Anti-CCRL2 ADC Drug-antibody ratio calculation**

| **Peak** | **Drug loading** | **Area %** | **DAR weight** |
| --- | --- | --- | --- |
| 2.6 | 0.0 | 0.0 | 0.0 |
| 6.3 | 2.0 | 19.1 | 0.4 |
| 7.5 | 4.0 | 29.9 | 1.2 |
| 8.9 | 6.0 | 35.4 | 2.1 |
|  |  | **Total DAR** | **3.7** |

**Supplementary Table 2.** Clinicopathologic features of MDS/AML patients and healthy donors for the *in vitro* analysis of anti-CCRL2 ADC activity

| **Characteristic** | ***TP53* mutated MDS/AML (N=10)** | ***TP53* wild-type MDS/AML (N=8)** | **Healthy donors (N=6)** |
| --- | --- | --- | --- |
| Age | 66 (27 – 73) | 57 (53 – 74) | 43 (24 – 50) |
| Gender  Females  Males | 3 (30%)  7 (70%) | 3 (37.5%)  5 (62.5%) | 2 (33%)  4 (67%) |
| Diagnosis  *TP53* mutated MDS/AML  Acute erythroid leukemia  De novo *TP53* wild-type AML  MDS-related *TP53* wild-type AML | 7 (70%)  3 (30%)  0 (0%)  0 (0%) | 0 (0%)  0 (0%)  4 (50%)  4 (50%) | N/A |
| Treatment-related | 4 (40%) | 1 (12.5%) | N/A |
| MDS-related changes | 7 (70%) | 4 (50%) | N/A |
| Blasts | 40% (10%-80%) | 40 (25%-60%) | N/A |
| Karyotype  Normal/Good  Intermediate (+8, del7)  Complex | 0 (0%)  0 (0%)  10 (100%) | 5 (62.5%)  3 (37.5%)  0 (0%) | N/A |

**Supplementary Table 3.** Variants and variant allele frequencies of patients with *TP53* mutated myeloid neoplasms for the *in vitro* analysis of anti-CCRL2 ADC activity

| **Variant** | **VAF** | **Other gene mutations** |
| --- | --- | --- |
| D281N | 60.6 | No |
| Y126H | 81.9 | *CBL*, *EZH2* |
| H178P | 82.4 | No |
| Y220C | 64.1 | No |
| D281N | 97.4 | *KIT* |
| P75fs | 54.4 | No |
| G245S | 67.5 | No |
| F270C | 89.9 | *NRAS* |
| R196P | 50.2 | No |
| V272M | 69.1 | *TET2* |

**Supplementary Table 4.** Clinical, pathological and molecular characteristics of multi-hit *TP53* mutated MDS/AML including in patient-derived xenograft studies

|  | Age at diagnosis | Sex | Pathologic diagnosis | Blasts% | Karyotype | Somatic mutations | *TP53* variant | Variant allele frequency | Mice engrafted | Treatment groups |
| --- | --- | --- | --- | --- | --- | --- | --- | --- | --- | --- |
| Pt 1 | 64 | M | AML with MDS-related changes | 30 | 43~47,XY,t(1;3;11)(p32;p23;p15),add(2)(p13),-3,add(3)(q21),add(4)(p14),del(4)(q25),dup(5)(q11.2q31),-7,  add(8)(p23),add(9)(p21),-12,?del(12)(q13q22),-17,?del(20)(q11.2),+1~4mar[cp20] | *DNMT3A, TP53, KMT2A* | V272M | 60.5 | 5/8 | 3 IgG2a  2 anti-CCRL2 |
| Pt 2 | 77 | M | AML with MDS-related changes | 40 | 44-47,XY,add(3)(q21),-5,add(5)(q31),add(6)(p21),+7,add(7)(q22),der(7;8)(p10;q10)x2,+8,i(8)(q10)x2,del(11)(q23q?25),del (12)(p11.2p12),add(16)(q24),der(16)t(16;21)(q12;q11.2),add(17)(q21),?19,del(20)(q11.2),-21,+r,+1-3mar[cp20] | *TP53* | F270C | 89.9 | 2/6 | 2 IgG2a  2 anti-CCRL2 |
| Pt 3 | 64 | M | AML with MDS-related changes/therapy related-myeloid neoplasm | 25 | 42,XY,der(5;17)(p10;q10),add(7)(q11.2),psu dic(22;14)(q13.3;p13),add(18)(p11.2),-19,-21[5]/43,sl,+mar[2]/46,XY[13] | *TP53* | Y220C | 64.1 | 2/6 | 1 IgG2a  1 anti-CCRL2 |
| Pt 4 | 59 | M | AML with MDS-related changes/therapy related-myeloid neoplasm | 22 | 43~49,XY,del(5)(q31q33),-7,-13,-18,+1~2r,+1~5mar[cp18] | *TP53* | H179R | 79.3 | 0/5 |  |
| Pt 5 | 27 | M | Acute erythroid leukemia | 40 | 46~47,XY,-5,add(9)(p24)x2,add(17)(p11.2),add(19)(p12),i(21)(q10),+1~2mar[8]/43~46,XY,-5,add(5)(q11.2),add(7)(q11.2),  -13,-13,-15,add(15)(p11.2),-16,add(17)(p11.2),-19,+21,add(21)(p11.2)x2,+1~4mar[cp9]/46,XY[3] | *TP53* | D281N | 60.6 | 0/5 |  |
